# Prostacyclin as a Negative Regulator of Angiogenesis in the Neurovasculature

**DOI:** 10.1101/2021.04.28.441854

**Authors:** Tasha R Womack, Jiabing Li, Pavel A Govyadinov, David Mayerich, Jason L Eriksen

## Abstract

In this study, multiple measures of angiogenic processes were assessed in murine brain endothelial (bEnd.3) cells after exposure to the stable prostacyclin analog, iloprost. Additionally, changes in the γ-secretase enzyme were evaluated after activation of prostacyclin signaling using PGI_2_ overexpressing mouse brain tissue and immunohistology studies in bEnd.3 cells. A three-dimensional assay of tube formation revealed that iloprost inhibits normal formation by significantly reduced tube lengths and vessel mesh area. The iloprost-mediated inhibition of tube-like structures was ameliorated by a specific IP-receptor antagonist, CAY10449. Reductions in wound healing were observed with iloprost application in a dose-dependent manner and this effect was reversed using CAY10449. Iloprost did not exhibit anti-proliferative effects in the bEnd.3 cells. When subjected to a Transwell assay to evaluate changes in trans-epithelial electrical resistance (TEER), bEnd.3 cells displayed reduced TEER values in the presence of iloprost an effect that lasted over prolonged periods (24 hours). Again, CAY10449 was able to reverse iloprost-mediated reductions in TEER value. Surprisingly, the adenylyl cyclase activator, forskolin, produced higher TEER values in the bEnd.3 cells over the same time. The TEER results suggest that iloprost may not activating the Gs protein of the IP receptor to increase cAMP levels given by the opposing results seen with iloprost and forskolin. In terms of γ-secretase expression, PGI_2_ overexpression in mice increased the expression of the APH-1α subunit in the hippocampus and cortex. In bEnd.3 cells, iloprost application slightly increased APH-1α subunit expression measured by western blot and interrupted the colocalization of Presenilin 1 and APH-1α subunits using immunohistochemistry. The results suggest that prostacyclin signaling within bEnd.3 cells is anti-angiogenic and further downstream events have effects on the expression and most likely the activity of the Aβ cleaving enzyme, γ-secretase.

## Introduction

Alzheimer’s disease (AD) is the leading form of dementia in people 65 years or older and is characterized by a progressive loss of thinking, behavioral and social skills. AD is a multifactorial disease involving extracellular deposition of amyloid plaques and intracellular neurofibrillary tangles leading to synaptic dysfunction, activation of microglia, decreased vascular tone and structure, and ultimately neuronal death. Epidemiological studies demonstrate considerable overlap between AD and cerebrovascular disease that causes an added or synergistic worsening of pathologies on cognitive decline [1, 2]. The two most common forms of vascular pathologies in AD are cerebral amyloid angiopathy and small vessel disease [3].

The first pathophysiological changes to occur in individuals with preclinical AD are of vascular origin including reduced cerebral blood flow and focal morphological changes in capillary networks [4, 5]. Our understanding of the intricate signaling mechanisms of angiogenesis have developed over the last 30 years with increased interest in angiogenesis dependent tumor research [6]. Angiogenesis, in physiological conditions, is a balance at the local level between endogenous stimulators and inhibitors of this process [6]. However, little is known about the intracellular processes that determine cerebrovascular damage in AD, and effective blood brain barrier protective substances for AD treatment have yet to be identified.

Epidemiological studies and clinical data suggest that nonsteroidal anti-inflammatory drugs (NSAIDs) are beneficial in the treatment and prevention of AD [7]. The protective effects of NSAIDs in AD have been attributed to the anti-inflammatory effects produced by cyclooxygenase-2 (COX-2) inhibition [8]. COX-2 plays a major role in inflammatory reactions in the peripheral tissues carried out by its metabolic products, prostaglandins (PGs), including PGE_2_, PGD_2_, PGF_2α_, TXA_2_, and PGI_2_. The vascular properties of these metabolic products have been investigated in models of ischemia and stroke. However, the pathophysiologic effects of PGI_2_ in the brain are not well studied. This report will continue investigations into the vascular effects of PGI_2_ within models of the cerebral vasculature.

Previous studies in our lab described exacerbated pathology and harmful cerebral vascular effects of prostacyclin overexpression in the APdE9 animal model of AD [9]. We found that upregulated prostacyclin expression significantly worsened cognitive abnormalities, accelerated amyloid pathology, and damaged the neurovasculature PGI_2_ overexpression selectively increased soluble amyloid β 42 production, total amyloid accumulation, and burden. PGI_2_ altered microvessel length and branching, and PGI_2_ expression in combination with amyloid was more detrimental than amyloid expression alone. Experimental studies suggest a role for PGI_2_ in the pathogenesis of AD through mechanisms relating to the γ-secretase cleavage enzyme of the amyloid precursor protein and recent reports have described that a key subunit of this enzyme, APH-1α, plays a significant role in both amyloid beta metabolism and angiogenesis [10].

In this study, we used functional, biochemical, and molecular approaches to investigate the effects of iloprost, a stable prostacyclin analog, on *in vitro* models of angiogenesis. While recognizing the potential role of γ-secretase in iloprost-mediated vascular effects, this study focuses on alterations to the enzyme’s subunits and their role in the inhibitory effects of iloprost during angiogenesis.

## Results

### Effects of iloprost on tube formation

Tubule formation is an integral process in angiogenesis. We utilized a three-dimensional Matrigel assay to assess the potential effects of iloprost application on brain vascular endothelial cells from mice (bEnd.3 cells, ATCC) after tube formation is induced. When bEnd.3 cells are applied to the thick Matrigel matrix, the basement membrane extract encourages the cells to morphologically differentiate into robust, pre-capillary “tubes” that is recognized as one of the initial steps in blood vessel formation [11] with peak tube formation occurring around 3-4 hours. Multiple tube characteristics were calculated using inverted-phase contrast microscopy, where two dimensional images of the top of the well were captured and analyzed with a 2D tracing software. As shown in Figure 1A, the total mesh area created by connected tubules was significantly reduced after iloprost treatment of 1 nM relative to the vehicle (P < 0.05). Also, tube length in the presence of 100 nM iloprost was significantly reduced to 4525 from the vehicle group at 5746 (P < 0.05; Figure 1B). These results demonstrate the potential of iloprost to inhibit tube-formation.

**Figure 1.**
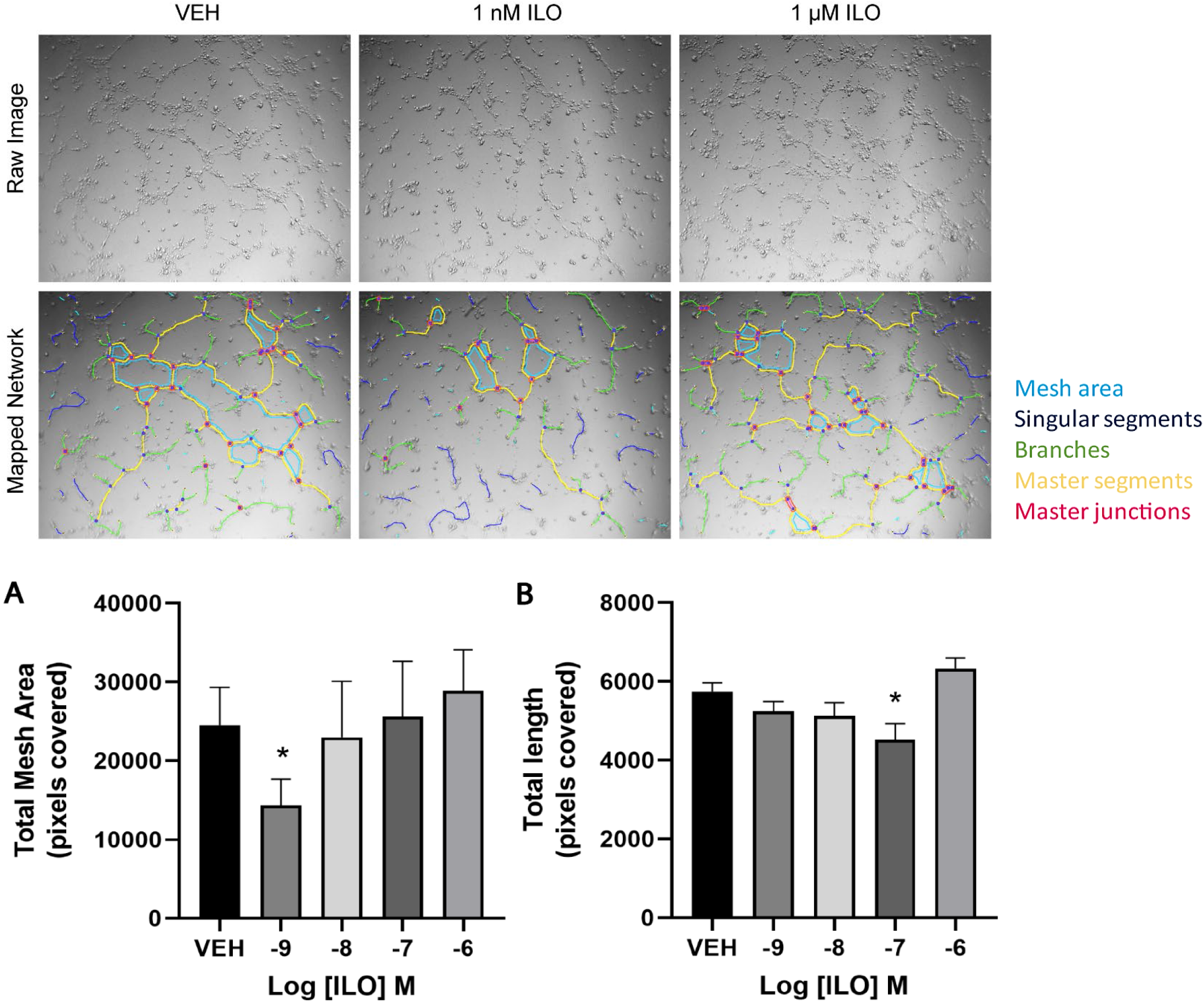
Prostacyclin analogue, Iloprost, inhibits pre-capillary tube formation in vitro. bEnd.3 cells were plated in triplicate at 50,000 cells per well in 48 well assay plates containing a 150uL layer of matrigel, a basement membrane extract. Peak tube formation is observed at 3 hours and phase contrast images were taken of each well in each quadrant. Representative 10X magnification images of five independent experiments. VEH – vehicle, ILO – iloprost. Reduced mesh area (A) and length (B) are observed in brain endothelial cells (bend.3) cells after 3 hours incubation with Iloprost. The data represent the means ± S.E.M. of five independent experiments. *p < 0.05 with respect to non-treated vehicle wells measured by unpaired t tests. VEH – vehicle, ILO – iloprost.

### Iloprost prevents wound healing

A well-established process of angiogenesis is cell migration after an injury or wound. To determine whether iloprost could inhibit cell migration we utilized a wound-healing assay to explore changes in migration following treatment with iloprost in the presence or absence of the IP-receptor antagonist, CAY10449. As shown in Figure 2A, identical injuries were made to confluent monolayers and images taken every 0, 4, 12 and 24 hours. The percentage of wound closure at 24 hours after wounding was about 50% in vehicle treated cells, whereas in 3 nM iloprost treated cells, the percentage decreased to 36.6% (P < 0.05, Figure 2B). Also, the wound healing rate at 24 hours was significantly reduced in the 3 nM iloprost group relative to the vehicle group (P < 0.05, Figure 2C). Addition of CAY10449 30 minutes prior to wounding and during the length of the assay was able to rescue iloprost-induced inhibition of cell migration into the wounded area (P < 0.05, Figure 3). These data show that iloprost reduces cell migration distance and this effect is mediated through the IP receptor as CAY10449 was able to reverse this inhibition.

**Figure 2.**
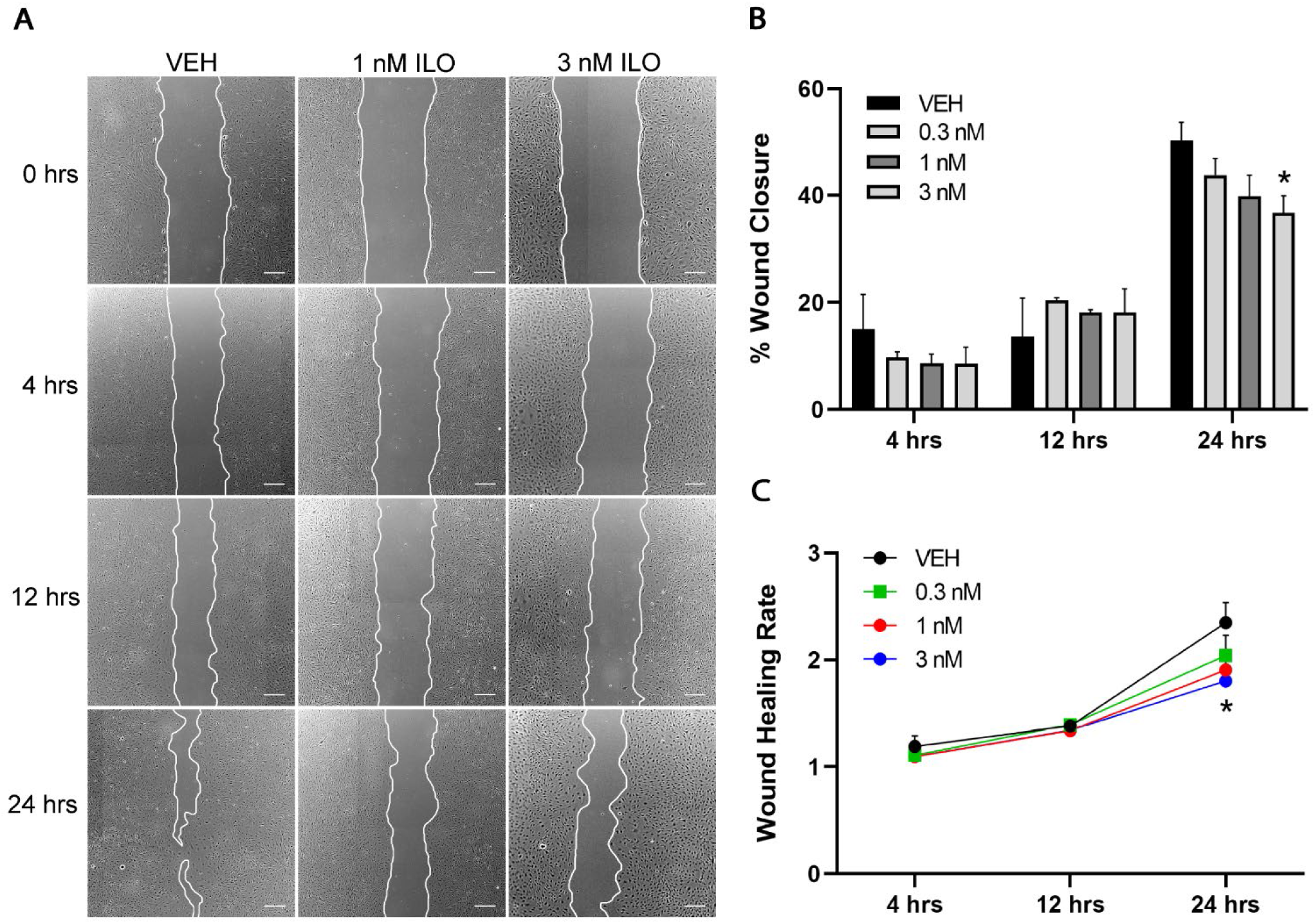
Delayed wound closure with iloprost application. (A) bEnd.3 cells were mechanically injured, and wound healing was followed over a 24-hour period (n=3-5, * *p < 0.05*). Representative images at 0, 4, 12 and 24 hours are presented. Scale bar = 200 μm. (B-C) Relative wound closure and healing rates over 24 hours. N = 3-6 independent experiments. Data are presented as the mean ± SEM. VEH – vehicle, ILO – iloprost, CAY – CAY10449.

**Figure 3.**
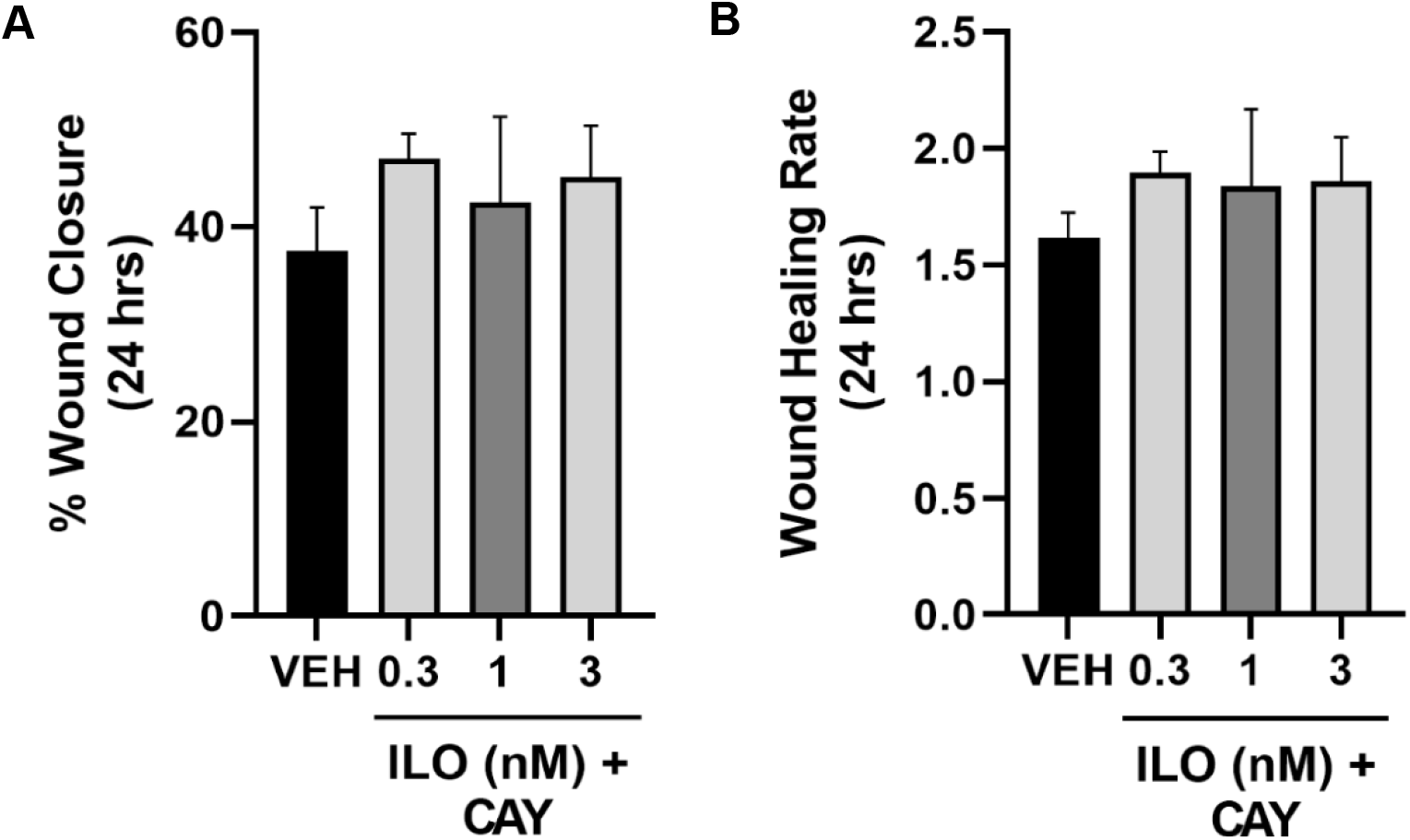
IP receptor-antagonist reversed iloprost-mediated delays in wound closure. Addition of CAY at a concentration of 10 nM for 30 minutes prior to iloprost application and during the 24-hour period prevents iloprost mediated delays in wound healing. N = 3 independent experiments. Data are presented as the mean ± SEM. VEH – vehicle, ILO – iloprost, CAY – CAY10449.

### Decreased TEER values in bend.3 cells after iloprost application

bEnd.3 cells grown on Transwell membranes were incubated with 1 nM iloprost, iloprost with 10 nM CAY10449 (Figure 4A). Iloprost application caused a significant decrease in TEER value at 3 hours and maintained this decreased TEER value over 12 hours (Figure 4B, p < 0.05). IP receptor inhibition with CAY10449 incubation in combination with iloprost ameliorated the decrease in TEER value to values to that of vehicle levels (Figure 4B). Notably, when CAY10449 was applied on its own TEER values significantly increased by 3 hours and this effect is sustained over the 24-hour period (Figure 4B, p < 0.05).

**Figure 4.**
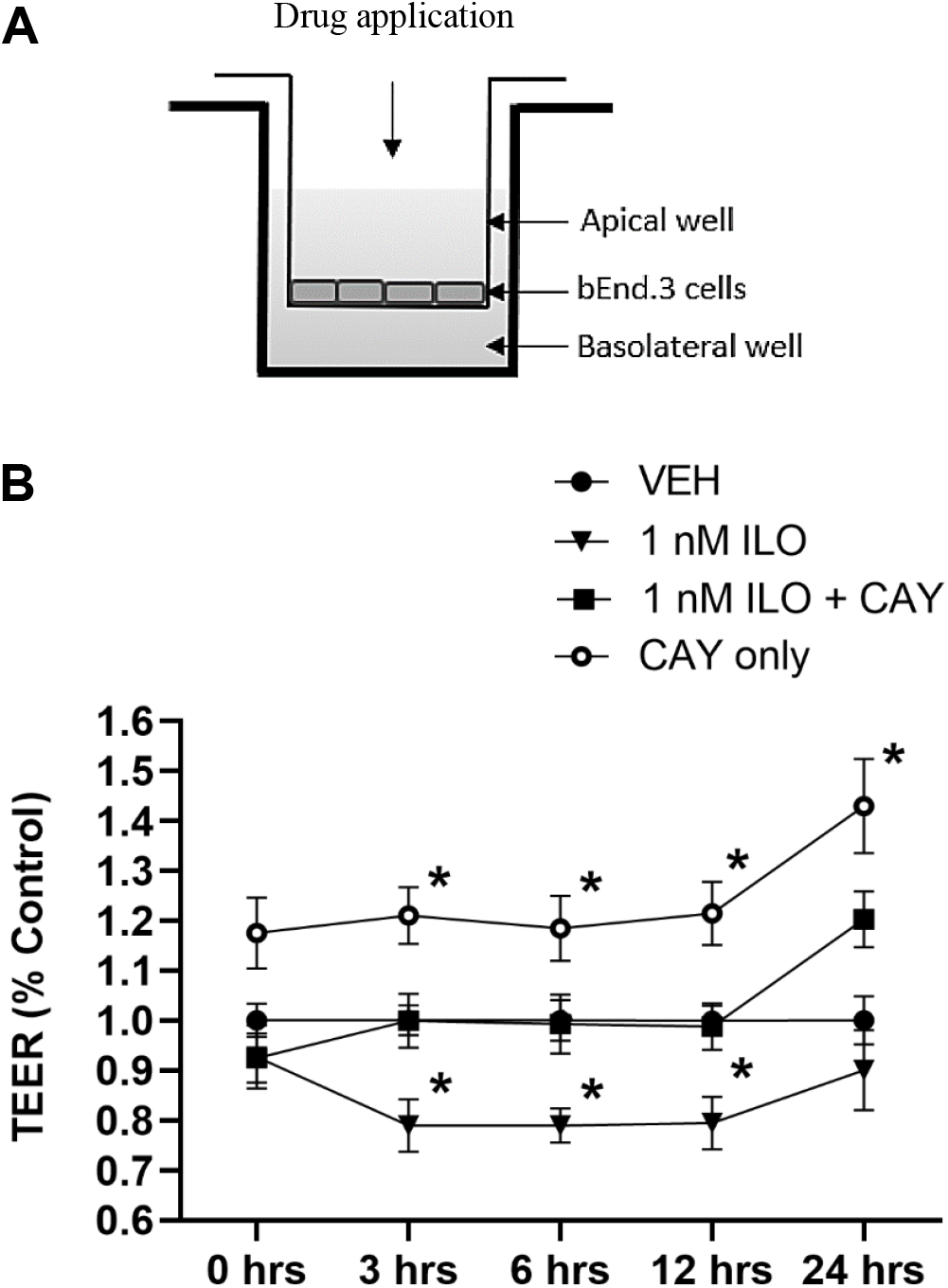
Decreased barrier integrity of bEnd.3 monolayers in presence of iloprost. (A) Schematic of the *in vitro* barrier model. (B) Relative TEER values evaluated for a 24-hour period on the 5^th^ day after seeding for iloprost, iloprost with 10 nM CAY10449 and 10 nM CAY10449 alone. CAY10449 was applied for a 30-minute pre-incubation before iloprost was added. N = 4 independent experiments. Data are presented as the mean ± SEM. * p < 0.05 with respect to the non-treated vehicle wells measured by one-way ANOVA. VEH – vehicle, ILO – iloprost.

To assess if the iloprost-mediated decrease in TEER value is related to elevated cAMP levels within the cells we decided to use the adenylyl cyclase activator, Forskolin, to simulate increased cAMP levels. Transwell membranes with bend.3 monolayers were treated with 10 μM Forskolin and TEER measured over a 24-hour period. Interestingly, the TEER value of bEnd.3 cells treated with Forskolin significantly increased after 3 hours and the effect became more significantly profound by hour 24 (Figure 5, p < 0.05).

**Figure 5.**
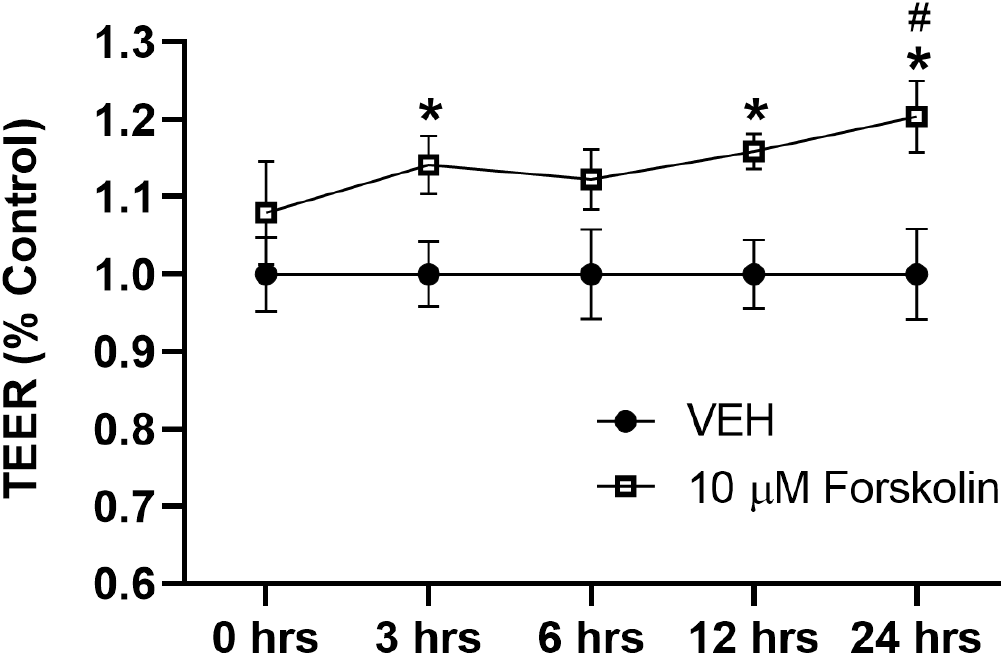
Forskolin increases barrier integrity of bEnd.3 monolayers measured by increased TEER values. TEER values evaluated for a 24-hour period on the 5^th^ day after seeding for adenylyl cyclase activator, forskolin. N = 3 independent experiments. Data are presented as the mean ± SEM. * p < 0.05 with respect to the non-treated vehicle wells measured by one-way ANOVA. # p < 0.05 with respect to 10 μM Forskolin at 0 hours measured by unpaired t tests. VEH – vehicle.

### Iloprost has no effect on cell proliferation in bEnd.3 cells

Another important process in the angiogenesis is cell proliferation. Prostacyclin signaling is known to exhibit anti-proliferate effects in VSMCs of the peripheral vascular system and in preclinical studies of HUVECs. Proliferation of bEnd.3 monolayers in the presence of iloprost for 3 hours was quantified using a Click-iT Edu incorporation assay. Representative fluorescent images of the EdU-positive cells are shown in Figure 19. The number of EdU-positive cells were unchanged in the presence of iloprost treatment at 0.01 nM – 100 nM (Figure 6).

**Figure 6.**
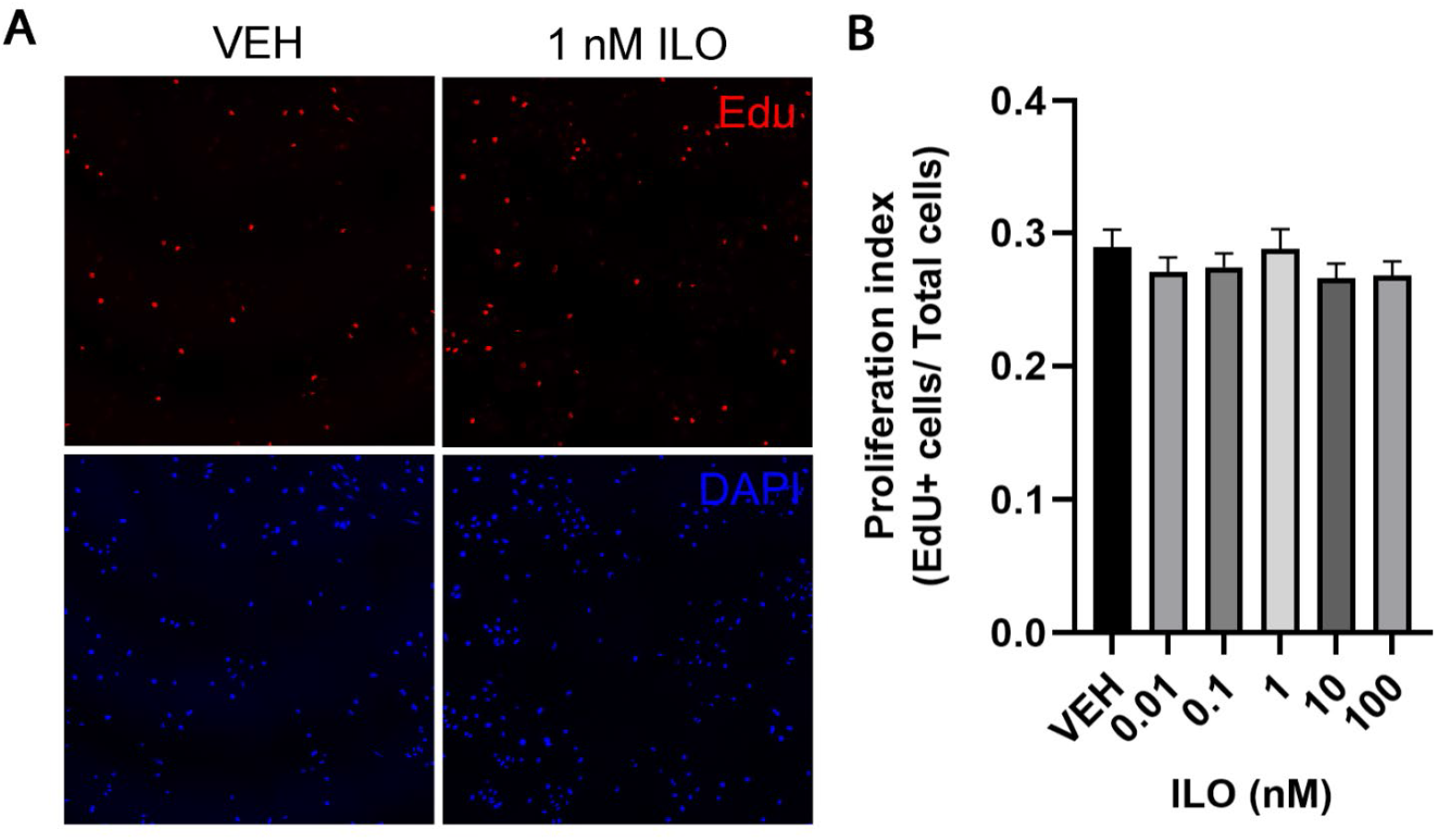
(A) Iloprost application has no effect on bEnd.3 cell proliferation. EdU (red) was used to detect the newly synthesized DNA of proliferating cells during the 3 hour iloprost treatment, and DAPI (blue) was used to measure total cell number. (B) The proliferation index of EdU positive cells to total cells was not affected by iloprost treatment N = 3. Data are presented as the mean ± SEM. One-way ANOVA. VEH – vehicle, ILO – iloprost.

### APH-1α is highly expressed in APdE9/CP-Tg transgenic mice

Previous evidence has indicated a pivotal role of the APH-1α subunit of the γ-secretase complex in the pathogenesis of AD [10, 12]. We evaluated the expression levels of APH-1α in an AD mouse model overexpressing prostacyclin synthase. As shown in Figure 7, immunostaining of APH-1α was markedly increased in the hippocampal neurons of both CP-Tg and CP-Tg/APdE9 mice and within neurons of the cerebral cortex in CP-Tg mice at 18-20 months of age when compared to age-matched controls (P < 0.05, Figure 7A&B).

**Figure 7.**
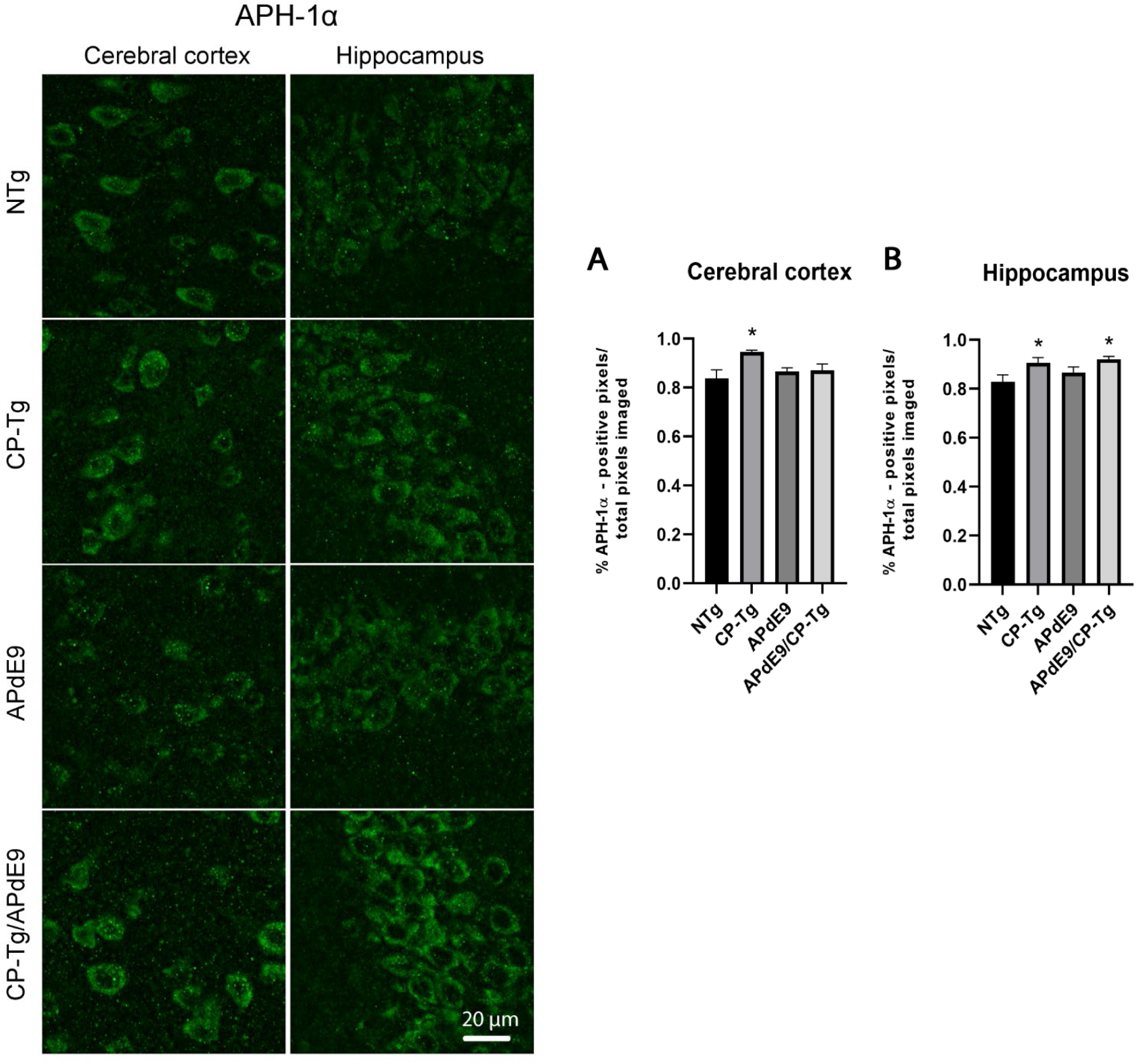
APH-1α is highly induced in mice expressing the CP-Tg genotype when compared to age-matched controls. Tissue blocks were created from brains excised and fixed in alcohol-free, Accustain followed by OCT embedding. The immunoreactivity of APH-1α was determined by immunofluorescence using an anti-APH-1α antibody on 60 μm free floating slices. Three randomly selected 100X z-stack images were captured in both the cerebral cortex and pyramidal cell layer of the hippocampus for all four genotypes. The immunoreactivity of APH-1α was quantified using intensity graphs of each 3D image stack separately for each region of interest, the cortex (A) or the hippocampus (B). N = 4-5 brains per group. Data are presented as the mean ± SEM. \**p < 0.05* with respect to the NTg controls measured by one-way ANOVA.

### Iloprost disrupts colocalization of Presenilin 1 and APH-1α subunits of the γ-secretase complex

Following our expression studies of the APH-1α subunit, we wanted to assess the interaction of this subunit to the catalytic subunit Presenilin 1 *in vitro* in response to iloprost stimulation. Monolayers of bEnd.3 cells were subjected to a 3-hour incubation with iloprost and subsequently fixed and stained for the APH-1α and Presenilin 1 subunits. High resolution images were captured and are represented in Figure 8. Colocalization analysis in ImageJ revealed that iloprost caused a significant interruption in the interaction of the two subunits measured by Manders Colocalization Coefficients. Specifically, the fraction of immunoreactive APH-1α overlapping or in direct contact with immunoreactive Presenilin 1 was significantly decreased after iloprost application (M1 coefficient) (P < 0.05, Figure 9). However, no change was detected in the percentage of immunoreactive Presenilin 1 overlapping immunoreactive APH-1α (M2 coefficient) (Figure 9).

**Figure 8.**
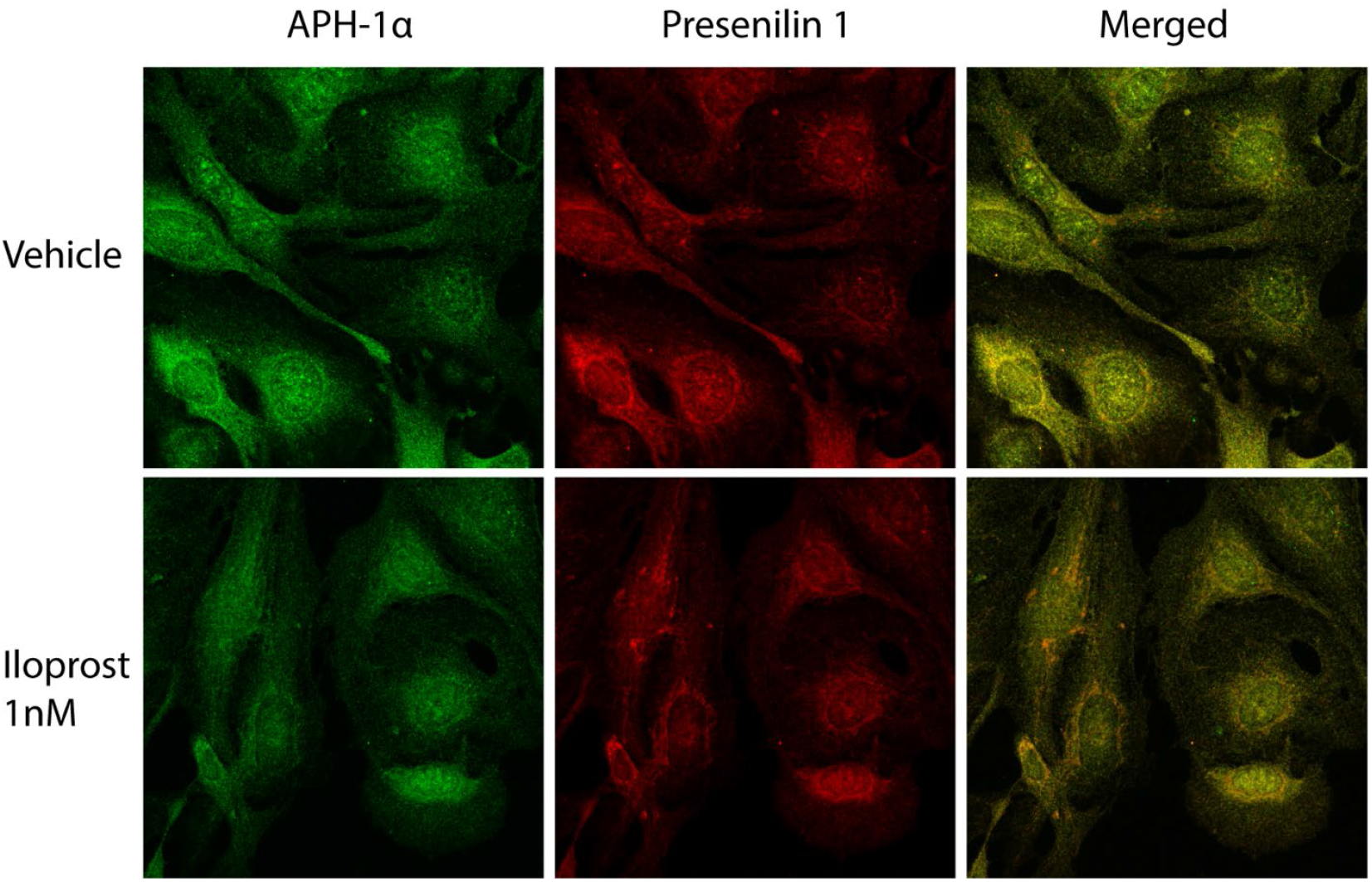
Representative images of disrupted localization of APH-1α with Presenilin 1 subunits after Iloprost treatment. bEend.3 cells were plated onto 12 mm coverslips given 2 days to grow and then treated with 1 nM of Iloprost for 3 hours. After incubation the cell monolayers were immediately fixed in 100% ice-cold methanol for 5 minutes at room temperature. Cells were permeabilized with 1X PBS containing 0.1% Triton X-100 for 10 minutes before a 1-hour protein block incubation in 1X TBST containing 5% donkey serum. Cells were then incubated with an anti-Presenilin 1 antibody at 1:200 overnight, biotinylated anti-rabbit secondary the next day for 1 hour and a streptavidin Dylight 649 fluorophore for 10 minutes. The cells were then given an avidin-biotin block. After washing the cells were then incubated in anti-APH-1α antibody at 1:200 overnight, a biotinylated anti-rabbit secondary the next day for 1 hour and a streptavidin Dylight 488 fluorophore for 10 minutes. Coverslips were then mounted onto slides and nine 100X z-stack images were captured for each of the control and Iloprost groups (three image stacks per coverslip with triplicate coverslips). N = 3 independent experiments.

**Figure 9.**
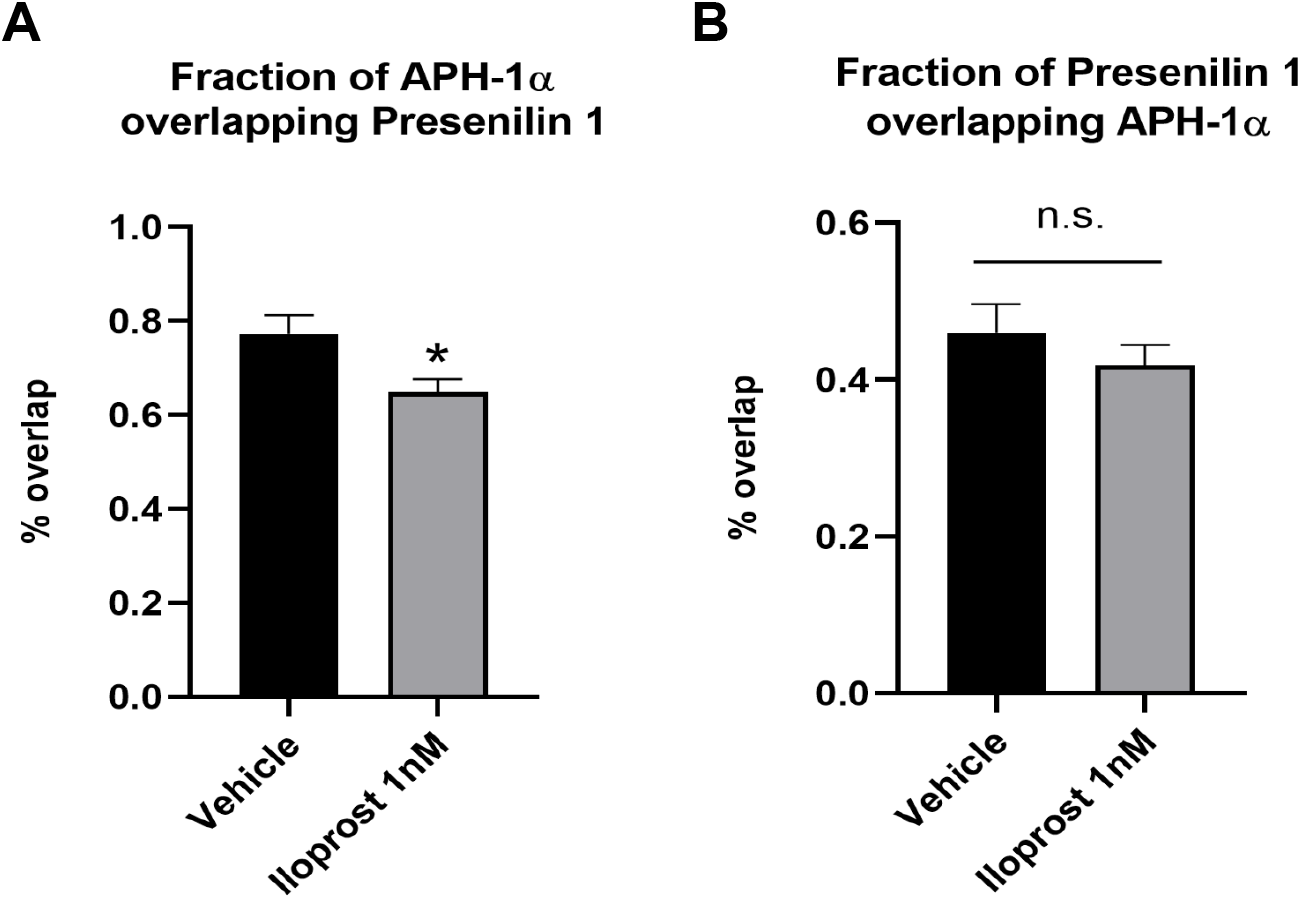
Colocalization analysis of Presenilin 1 and APH-1α subunits of γ-secretase after iloprost treatment. Manders colocalization coefficients (MCCs) quantified from dual immunofluorescent images of APH-1α- and Presenilin 1-positive bEnd.3 monolayers. The M1 coefficient signifies the fraction of immunodetectable APH-1α overlapping Presenilin 1 (A). Whereas the M2 coefficient signifies the fraction of immunodetectable Presenilin 1 overlapping APH-1α (B). N = 3 independent experiments. Data are presented as the mean ± SEM. * *p < 0.05* with respect to vehicle measured by unpaired t tests. n.s. – not significant.

### Western blot analysis of γ-secretase subunits after iloprost application in bEnd.3 cells and in APdE9/CP-Tg mice

Mouse bend.3 cells were treated with increasing concentrations of Iloprost for 3 hours before extracting protein for western blotting. One-way ANOVAs were used to analyze differences in protein levels in response to increasing doses of iloprost. No significant differences were found in the APH-1α, Presenilin 1, or Nicastrin subunits of the γ-secretase enzyme at any dose (Figures 10). Iloprost application did not appear to significantly influence the levels of Notch 1 and Notch 3 cleaved intracellular signaling domains (Figure 11).

**Figure 10.**
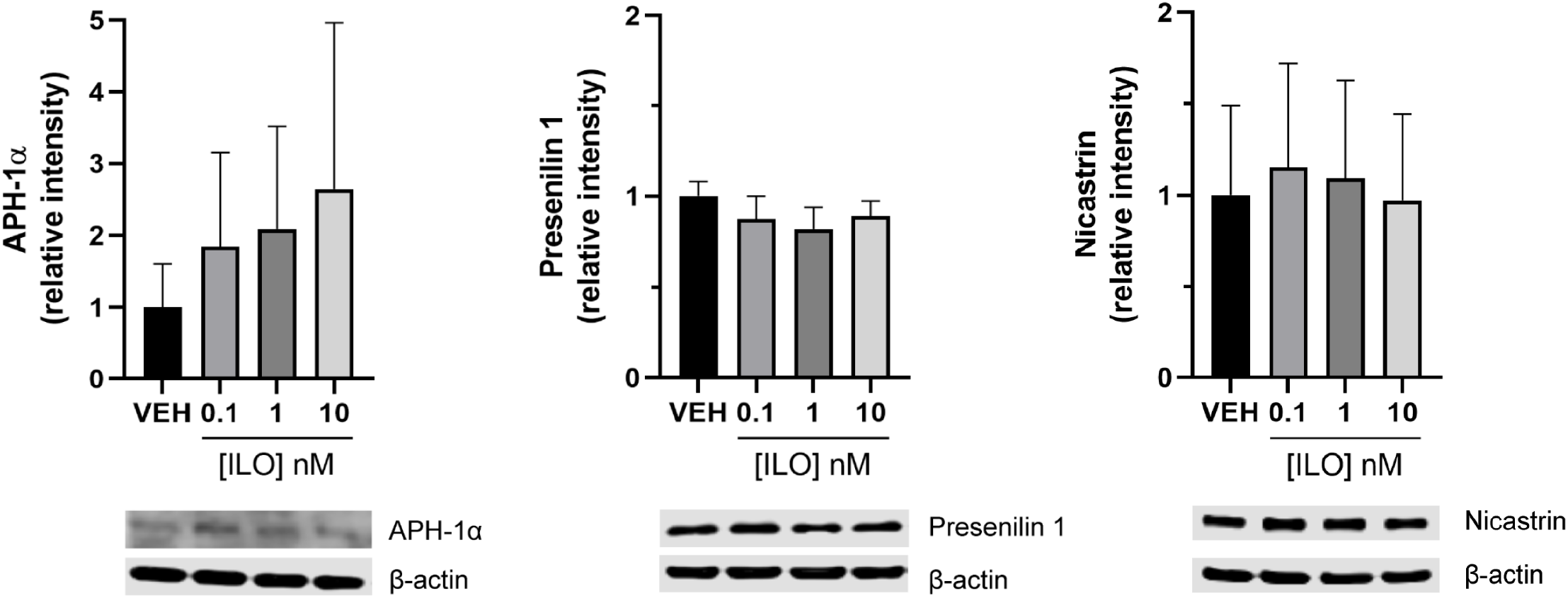
Western blot analysis of APH-1α and Presenilin 1 protein levels in bEnd.3 cells after a 3-hour incubation with iloprost. Iloprost did not significantly alter APH-1α, Presenilin 1, or Nicastrin protein levels in bEnd.3 cells. Total β-actin levels were used to assess consistent lane loading. N = 3. Data are presented as the mean ± SEM. One-way-ANOVA.

**Figure 11.**
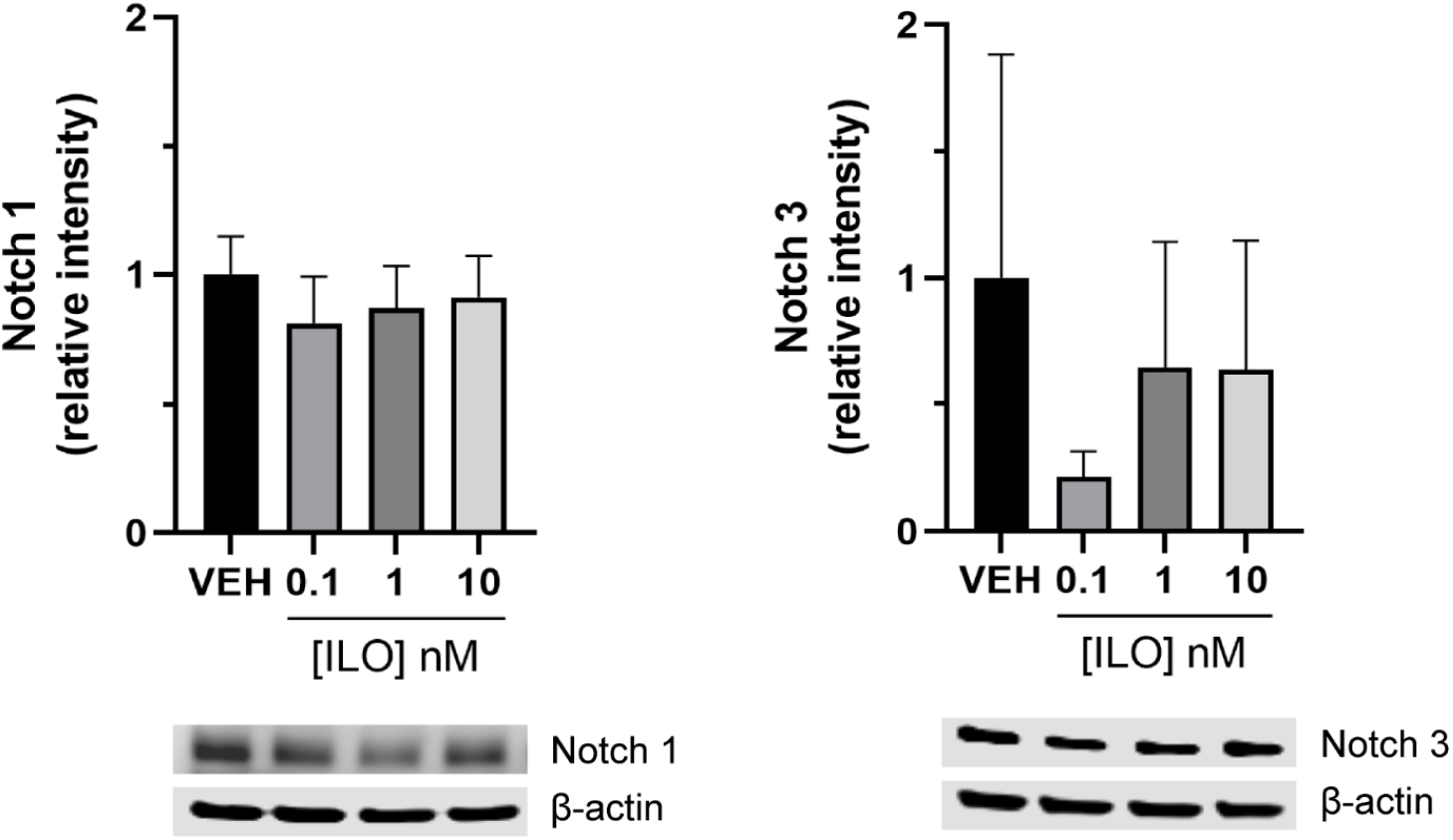
Western blot analysis of Notch 1 and Notch 3 protein levels in bEnd.3 cells after a 3-hour incubation with iloprost. Iloprost did not change Notch 1 or Notch 3 protein levels in bEnd.3 cells. Total β-actin levels were used to assess consistent lane loading. N = 3-5. Data are presented as the mean ± SEM. One way ANOVA.

Brain homogenates of prostacyclin overexpressing mice and their respective controls (NTg, CP-Tg, and APdE9 mice) were lysed in RIPA buffer and assessed using western blotting. One-way ANOVAs were used to analyze differences between the four groups. Western blot analysis revealed no significant changes in the Presenilin 1 protein levels between the groups (Figure 12). The APdE9 and APdE9/CP-Tg brain homogenates exhibited significant increases in mature nicastrin with no changes in the immature form of nicastrin (Figure 12, p < 0.05). No changes in Notch 1 intracellular signaling domain were observed between the different groups (Figure 13).

**Figure 12.**
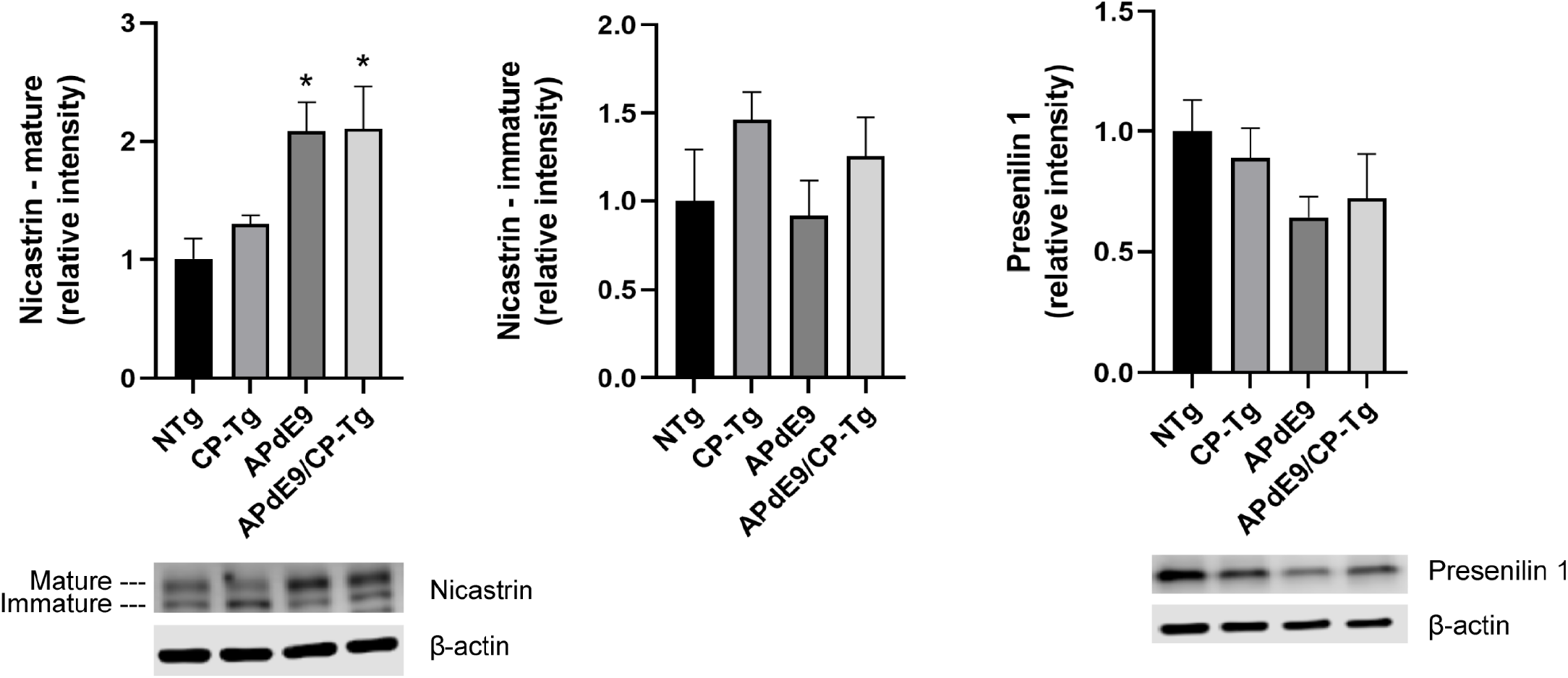
Western blot analysis of Nicastrin and Presenilin 1 protein levels in APdE9/CP-Tg mice. APdE9 mice models exhibited increased protein levels of mature Nicastrin. However, protein levels of immature Nicastrin remain unchanged among the four mouse genotypes. Neither genotype displayed a significant change in Presenilin 1 protein levels. Total β-actin levels were used to assess consistent lane loading. N = 4-5 mice per group. Data are presented as the mean ± SEM. One-way ANOVA.

**Figure 13.**
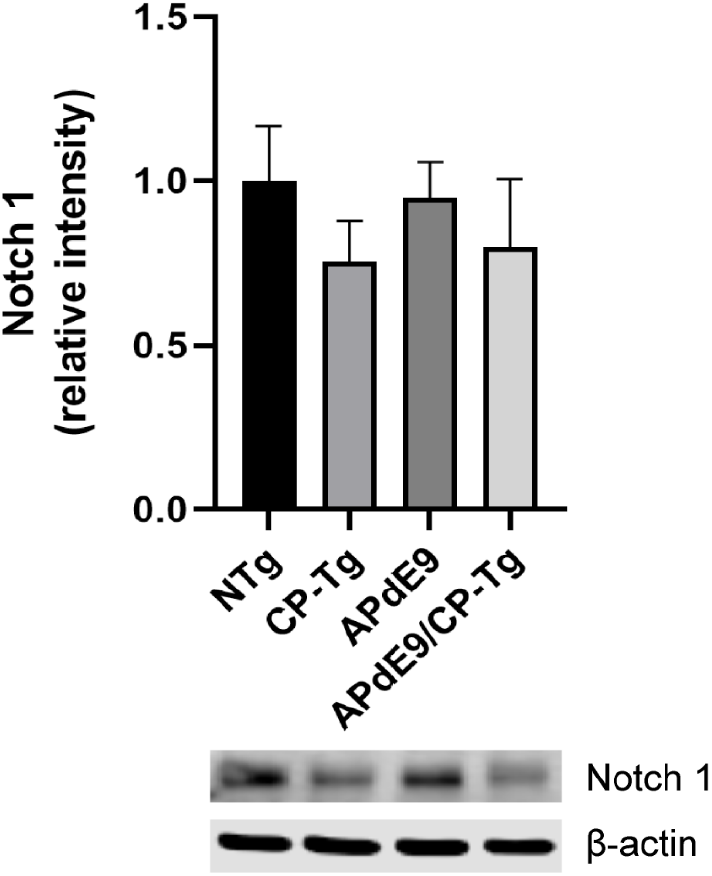
Western blot analysis of Notch 1 protein levels in APdE9/CP-Tg mice. None of the genotypes had a significant change in Notch 1 protein levels. Total β-actin levels were used to assess consistent lane loading. N = 4-5 mice per group. Data are presented as the mean ± SEM. One-way ANOVA.

## Discussion

In this study we evaluated the response of bEnd.3 cells to exogenous application of the vasodilator and platelet inhibitor, iloprost, by the quantification of capillary-like tube formation, wound repair, proliferation, and permeability. Our present findings confirm that stimulation of brain endothelial cells by the prostacyclin analogue, iloprost, restricts multiple measures of angiogenesis. Treatment with iloprost *in vitro* reduced pre-capillary tube formation and suspended wound healing abilities but had no effect on short term proliferation. Monolayer integrity measured by TEER displayed reduced resistance values in the presence of iloprost; however, this effect appears to be cAMP dependent as the adenyl cyclase activator, forskolin, showed increased TEER values within the same time frame. These data suggest that IP receptor activation plays a significant physiologic function in disrupting angiogenic processes and increasing vascular permeability. In this context, we describe a new role for prostacyclin in the regulation of vascular homeostasis in models of angiogenesis within the cerebral vasculature.

Prostacyclin is released from the endothelium as a vasoactive molecule after activation of COX-1 by physical or chemical stimuli. Hallmarks of AD include the deposition of amyloid plaques and neurofibrillary tangles alongside chronic inflammation caused by impaired clearance of these aggregated proteins. In response, the vascular endothelium produces and releases prostacyclin to serve as a vasodilator to increase blood flow and migration of leukocytes to the site of damage as well as promote oedema in the surrounding tissue [13]. Alongside these properties, prostacyclin has been recognized for its participation in regenerative processes of the cardiovascular system after endothelium injuries in events such as stroke, pulmonary hypertension and pulmonary fibrosis [14– 17]. The implication of these prostacyclin-mediated effects to the brain vasculature during pathogenesis of AD are less studied.

Prostacyclin is well documented for maintaining vascular homeostasis and inhibiting smooth muscle cell remodeling [13, 15, 18, 19]. Herein we demonstrate an inhibitory effect of iloprost on brain endothelial cell differentiation into tubules and endothelial cell migration after wound infliction. Limited comparisons are available, yet one study found that iloprost applied to an *in vivo* angiogenesis model of chick chorioallantoic membranes and an *in vitro* tube formation assay of human umbilical vascular endothelial cells (HUVECs), no significant changes in vessel growth or tube formation were observed, respectively [20]. Conversely, HUVECs treated with the IP receptor agonist, taprostene, exhibited increased tube formation *in vitro* when applied to basement membrane extract and increased cell migration measured by a matrigel droplet migration assay [21]. Notably, when higher concentrations of taprostene were given migration reduced to only 26% of the control at concentrations up to 25 μM and tube length fell to 29% of control at concentrations up to 25 μM [21]. They concluded that this effect was due to the deactivation, internalization and subsequent phosphorylation of the IP receptor as a G-protein coupled receptor by intentionally causing desensitization with high concentrations [21].

To the best of our knowledge this is the first time that a prostacyclin analogue has proven to be anti angiogenic. This suggests that the endothelium in the brain reacts is ways yet to be understood, a new function of prostacyclin in the context of the cerebrovasculature. A similar behavior is observed between the pulmonary endothelium and the systemic endothelium when reductions in oxygen concentration can induce opposite responses [22]. For example, hypoxia in the pulmonary endothelium leads to vasoconstriction, whereas hypoxia in the systemic arterial endothelium causes a vasodilatory response [22]. Farber concludes that endothelial cells are the initial cell in detecting alterations in blood flow and oxygen and depending on the tissue needs could likely have varying oxygen-sensing mechanisms activated in response to low concentrations [22]. As angiogenesis is a process sensitive to changes in blood flow and oxygen concentrations, there is basis for varying angiogenic responses between endothelial cells of two different vascular beds.

We found that acute incubations of iloprost also decreased brain endothelial cell resistance indicating an increase in vascular permeability. TEER assays are a widely accepted parameter to assess barrier integrity as it replicates tight junction dynamics that can be measured in various cell culture models such as monolayers or cocultures. Changes in TEER value can then be monitored in real time after application of drugs or chemicals. Previous literature is complementary to our findings. Early reports found that intradermal injections in mice with exogenous prostacyclin combined with nitric oxide donors or bradykinin synergistically increased vascular permeability, but not when administered alone [23, 24]. This indicates an interaction between prostacyclin and another vasoactive substance is necessary to induce vascular permeability *in vivo*, whereas we have shown PGI_2_ is capable of this on its own *in vitro*, specifically in endothelial cells. A more recent report showed that iloprost applied at 1 μM prevented LPC-mediated vascular barrier disruption in an *in vitro* BBB model (consisting of rat brain endothelial cells, pericytes and astrocytes) [25]. However, when iloprost was applied alone to the BBB model or monocultures of rat brain endothelial cells they found no significant changes in the TEER value [25]. An important difference to note from this study is the high dose of 1 μM used that is likely to induce greater desensitization of IP receptors given that the recorded inhibition constant (k_i_) of iloprost is 3-10 nM [26, 27].

Our previous work in mice indicates that amyloid burden is directly affected by prostacyclin expression. We found selective increases of Aβ42 in the soluble pool from AD mice overexpressing PGI_2_, suggesting a role for PGI_2_ in bias γ-secretase processing. To further investigate a functional relationship between PGI_2_ and γ-secretase we assessed changes in the enzyme’s subunits when exposed to PGI_2_. We found significantly higher levels of immunoreactive APH-1α in brain slices of both PGI_2_ overexpression models, the single transgenic and the double transgenic with AD. Notably, although the production of the APH-1α subunit was influenced by PGI_2_ overexpression alone we do not see a congruent increase in Aβ40 or Aβ42 levels in whole brain homogenates. These results suggest that PGI_2_-mediated increases in γ-secretase subunits do not on its own lead to a pathogenic amyloid phenotype, it is only the combination of PGI_2_ and an AD genotype that causes synergistic increases in Aβ42.

The results of IP receptor activation *in vitro* demonstrated a reduced colocalization of the Presenilin 1 and APH-1α subunits from the position of APH-1α. Two interpretations can be made of these results considering we know PGI_2_ has repeatedly be shown to increase γ-secretase cleaving activity. First, reduced colocalization may be due to an increase in the production of the APH-1α subunit and perhaps these newly synthesized proteins have yet to bind to a γ secretase complex within the endoplasmic reticulum (Lee et al., 2002 J Biol Chem). Second, increased activity of the enzyme may lead to a recycling of whole complexes that would causes the subunits to be separated for a time. Although, these interpretations are limited due to the single time point captured and more delineating results could have been observed if images were taken over time. These findings are supported by recent work on prostacyclin signaling influencing APP processing mechanisms. Agonist activation of the IP receptor was found to increase the expression of the amyloid precursor protein, resulting in the increased production of Aβ [28]. Another report suggests that downstream activation of the PKA/CREB pathway after prostacyclin injections induces upregulation of the APH-1α subunit thus increasing γ-secretase-dependent cleavage that leads to an accumulation of Aβ c-terminal fragments in an APP/PS1 transgenic mouse model [10].

The γ-secretase complex is formed through a concerted assembly of four subunits involving trafficking through the endoplasmic reticulum to the Golgi in a process that matures the enzyme to then be recycled through lysosomes (~95%) or transported to the cellular membrane for substrate cleavage (~5%) [29]. Each of the four subunits provides an integral role for which all subunits are required for γ-secretase activity [30]. In response to IP receptor activation, no significant changes in γ-secretase subunit protein levels were observed in bEnd.3 cells. However, the levels of APH-1α did show an upward trend to increasing doses of iloprost which would be consistent with previous results and literature describing PGI2 as a stimulator of APH-1α production [10]. A limitation of this study would be that the bEnd.3 cells are native cells with no additional AD related mutations which may lead to reduced or limited γ-secretase activity possibly explaining the absence of robust changes in subunit production. Another possible limitation as to why subunit levels remained unchanged is that they were assessed in non-neuronal, vascular cells, while the expression of γ-secretase is ubiquitous across cell types the extent of expression and protein regulatory mechanisms between neuronal and non-neuronal γ secretase may differ [31]. No changes in the Notch 1 or Notch 3 cleavage substrates of γ-secretase involved in angiogenesis were observed as well. A possible explanation as to why no changes were observed in cleaved Notch products could be that protein levels were probed from cell lysates of monolayers. This is significant because endothelial cells require external stimulus to initiate angiogenic processes (i.e. tube formation) as part of a differentiation process and monolayers would not express the same profile of proteins as that of endothelial cells in capillary formation. To elucidate the impact of PGI_2_-mediated changes to Notch receptor signaling through altered γ-secretase activity, future studies will need to focus on assessing Notch in endothelial cells *in vivo* or in actively differentiating endothelial cells.

We also assessed protein levels of two subunits, Presenilin 1 and Nicastrin, in whole brain homogenates of APdE9 mice overexpressing prostacyclin. We observed a downward trend in Presenilin 1 expression only in the APdE9 mice, but this effect was not significant. Whereas levels of mature Nicastrin were increased in both APdE9 mice and APdE9/CP-Tg mice. However, the increase in mature Nicastrin levels were similar for both mouse groups suggesting that this increase is an amyloid-mediated effect alone and prostacyclin overexpression was not a contributing factor. These results are congruent with previous reports that levels of mature Nicastrin are higher than immature levels in the presence of Presenilin heterodimers due to the formation of more stable γ-secretase complexes [29]. When observing downstream cleavage of Notch 1 receptors, we observed no change in cleaved Notch 1 domains. These results do not rule out the possibility that γ-secretase activity was increased in prostacyclin overexpressing mice. The use of whole brain homogenates for western blot analysis likely masked possible changes in protein levels within specific brain regions that were not robust enough to show in the whole homogenate. A limitation to this study is that the whole brain homogenates also combined all cerebral cell types so changes in neuronal vs non-neuronal cell types would be difficult to determine.

## Conclusions

This study evaluated the effects prostacyclin signaling on angiogenesis measures in *in vitro* models of murine brain endothelial cells. Our previous work in a mouse model of AD showed prostacyclin expression negatively impacted multiple aspects of vascular health and this process was exacerbated with Aβ pathology. Given that PGI_2_ has reportedly cardioprotective mechanisms in peripheral tissues, this study aimed at delineating the cellular processes leading to the detrimental vascular effects in the cerebrovasculature. We found that the stable prostacyclin analog, iloprost, inhibited various angiogenesis processes, including tubule formation, tubule length, and wound healing. Tight junction integrity was disrupted measured by reduced TEER values in response to iloprost, but the proliferation of bEnd.3 cells was unchanged. The use of an IP receptor antagonist was able to reverse the iloprost-mediated reductions and increase tube formation and TEER. Chronic vascular dysfunctions most likely play a synergistic role in neurodegenerative changes to advance cognitive impairment in individuals with AD [32]. Understanding the underlying mechanisms regulating angiogenic remodeling mechanisms within the cerebral vasculature cells would provide novel therapeutic targets for vascular damage in AD. We conclude that cerebrovascular remodeling by the vasoactive signaling molecule, iloprost, is a potential contributor to AD cerebrovascular dysfunctions.

Prostacyclin-mediated effects to the γ-secretase complex components and its downstream cleavage substrate, Notch 1, were also assessed. In mouse models of PGI2 overexpression, we found increased APH-1α subunit production with previous reports of PGI2 regulating expression of the APH-1α/β subunits [10]. In bEnd.3 cells, iloprost application disrupted the interaction of the Presenilin 1 and APH-1α subunits suggesting altered γ-secretase activity. However, no significant iloprost or PGI_2_-induced changes in γ-secretase and Notch were observed via western blot analysis. These observations further suggest a new relationship between PGI_2_ signaling and γ-secretase dynamics that potentially contribute to the pathogenesis of AD.

## Materials and Methods

### Brain endothelial (bEnd.3) cell culture

bEnd.3 cells were purchased from ATCC (CRL-2299) and cultured according to manufacturer’s recommendation using complete growth medium (10% fetal bovine serum in Dulbecco’s Modified Eagle’s Medium (DMEM)). Cells were used between passage 29 and 32 for all assays. Conditioned media for the tube formation assay was obtained by plating cells, allowing two days for the cells to recover and reach ~30% confluence and then replacing the media with 1% FBS media for 24 hours. The conditioned media was collected at 24 hours and immediately frozen at −80°C until the time of the assay.

### Endothelial capillary-like tube formation assay

Matrigel® basement membrane extract (0.15 mL, Cat. No. 356234, Corning) was added to each well of 48-well culture plates (Greiner Bio-One) and incubated for 30 minutes at 37°C creating a thick layer. Iloprost dilutions were prepared in 0.2 mL of 1% FBS media and applied to the respective wells. bEnd.3 cells were lifted by 0.25% trypsinization and resuspended in conditioned media (prepared as described above). From the cell suspension, 50,000 cells were applied to each well in triplicate for each treatment group, gently pipette mixed in the well and incubated for 3 hours at 37°C. Peak tube formation was determined to be 3 hours for bEnd.3 cells (data not shown). At 3 hours, the endothelial tubes were digitally imaged using a phase-contrast microscope (Leica Microsystems). The relative mesh area and length were determined by a 2D tracing software (ImageJ). Data are presented as relative average ± SEM of mesh area or total length, compared to controls, obtained from five independent experiments.

### TEER measurements

Before seeding, 12-well Transwell™ tissue culture inserts (Corning Costar Inc., Corning, NY) were coated with collagen, type I from rat tail (Cat. No. C3867, Millipore Sigma) at 10 μg/cm^2^ for two hours in a 37°C incubator. bEnd.3 cells were cultured to confluence for 5 days on the collagen-coated inserts at a seeding density of 1.5 × 10^5^/cm^2^. An EVOM voltohmmeter (World Precision Instruments Inc., Sarasota, FL) was used to measure TEER and experiments were performed once the monolayers reached a TEER of >30 ohm-cm^2^. Initial TEER measurements were taken in complete growth media before replacing the media in the apical chamber with iloprost diluted in 1% FBS media and the basolateral chambers with fresh 1% FBS media. TEER of blank inserts were subtracted from the measured TEER of test wells. TEER was determined in duplicate and obtained from four independent experiments.

### Wound healing assay

Cells were seeded at 50,000 cells/well into 48-well plates and given 5 days to reach confluence. Scratches were made with a plastic 1-200 μl pipette tip across the diameter of each well. Cell debris was removed with a quick wash of complete media before iloprost dilutions in 1% FBS media were added. Images of the wound area were collected using a Celigo^TM^ Imaging Cytometer (Nexcelom, Lawrence, MA). Quantitative analysis of the scratch area was performed using ImageJ (*MRI Wound Healing Tool* by Nathalie Cahuzac, Virginie Georget, and Volker Baecker, http://dev.mri.cnrs.fr/projects/imagej-macros/wiki/Wound_Healing_Tool.)

### EdU incorporation assay

Proliferating cells were detected using the Click-iT Plus EdU Alexa Fluor 594 Imaging Kit (cat. No. C10639, Life Technologies). Briefly, bEnd.3 cells were plated at 20,000 cells/well into 24 well plates containing sterilized 12mm round coverslips at the bottom of each well. Cells were given 24 hours to recover before EdU incorporation. The next day, half of the media was replaced with fresh 1% FBS media containing 20 μM EdU and increasing concentrations of iloprost and incubated for another 3 hours. Cells were fixed with ice-cold methanol, permeabilized with 0.5% Triton X-100 and stained with the reaction cocktail prepared according to the manufacturer’s instructions. Coverslips were removed from the wells and mounted onto microscope slides using Fluorogel II with DAPI (cat. No. 17985-50, Electron Microscopy Sciences). Coverslips were imaged with confocal microscopy and the proliferation index was evaluated by dividing total cell number by EdU-positive cells.

### Western blotting

Cultures of bEnd.3 cells at 80% confluency were incubated for 3 hours with Iloprost in 1% FBS media. After treatment, the cells were rinsed with cold 1X PBS before being scraped and homogenized in 0.1 mL of RIPA buffer containing 1% protease inhibitor cocktail (Sigma). Rabbit antibodies against Presenilin 1, APH-1α, Nicastrin and Notch 1 intracellular domain were obtained from Cell Signaling Technology. Rat antibody against Notch 3 intracellular domain was obtained from Biolegend. Blots were probed with a mouse anti-actin (ThermoFisher) antibody as loading controls. Protein was loaded at 15 ug/well and electrophoresis was performed for 1.5 hours at 120V using TGX premade gels and BioRad Mini PROTEAN Tetra electrophoresis cells. The separated protein was then wet transferred onto PVDF membrane for 1 hour at 100V. Blots were then blocked in TBST with 0.5% non-fat dried milk for 1 hour at room temperature and further incubated with primary antibodies between 1:100 and 1:1000 dilutions overnight at 4°C. The blots were washed and then incubated in HRP-conjugated secondary antibodies respective to the species of the primary for 1 hour at room temperature. Chemiluminescence was detected using SuperSignal West Femto substrate (ThermoFisher) and a C-DiGit western blot scanner (Li-cor).

### Immunofluorescent staining

In vitro immunostaining was carried out on bEnd.3 cultures plated onto coverslips. At 50% confluency the cells were treated with Iloprost in 1% FBS medium for 3 hours at 37°C. After drug incubation the cells were rinsed in cold 1X PBS and then fixed with ice-cold 100% methanol for 5 minutes at room temperature. Cells were then washed with TBST, permeabilized with 0.1% Triton X-100 for 10 minutes at room temperature and blocked in TBST containing %5 normal goat serum for 30 minutes. Cells were then incubated with a rabbit anti-Presenilin 1 antibody overnight at 4°C, washed with TBST, and incubated in buffer containing biotinylated anti-rabbit secondary antibody for 1 hour at room temperature. The cells were then washed three times before incubation in DyLight fluorophore solution for 10 minutes. The cells on coverslips were then blocked using an avidin-biotin blocking kit (Thermo) before continuing to the next primary antibody. Cells were then incubated with anti APH1α antibody, biotinylated anti-rabbit secondary antibody and DyLight fluorophore as described previously before being mounted onto slides for confocal imaging.

### Immunostaining in brain slices

Brain slices were obtained from our lab as previously described (see section 2.2.9). Briefly, the APdE9/CP-Tg mouse model was developed by crossing CP-Tg mice, that express a hybrid enzyme complex linking COX-1 to PGIS by a 10 amino acid linker (COX-1-10aa-PGIS), with APdE9 mice, a model expressing the human APP Swedish mutation (AP) and a variant of the exon-9-deleted presenilin-1 gene (dE9). Confirmation of genotype was determined by PCR analysis and heterozygous male and female mice were used for this study. Brain were acquired by decapitation and post fixation in the formalin free fixative, Accustain. Slices were obtained by paraffin embedding the brains and cutting 60 μM thick sections on a microtome. The free-floating coronal brain slices were then subjected to the same dual staining protocol as the bEnd.3 cells described above.

### Confocal microscopy and statistical analysis

For co-localization analysis in bend.3 cells, coverslips were then mounted onto slides and nine randomly selected 100X z-stack images were captured for each of the control and Iloprost groups (three image stacks per coverslip with triplicate coverslips) for three independent experiments. Co-localization of Presenilin 1 and APH-1α was performed using the ImageJ JACoP plugin as previously described [33]. All z-stacks were processed with background subtraction and analyzed using automatic thresholding. An unpaired t-test was used to analyze the bend.3 cell co localization data. For quantification of APH-1α immunoreactivity, three randomly selected 100X z-stack images were captured in the both the cerebral cortex and pyramidal cell layer of the hippocampus for all four genotypes. Raw images were run through a cell segmentation software [34] to create a binary mask of each image that segmented the cells from the rest of the image. A manually chosen threshold based on the background subtraction of the NTg images was applied evenly across the data set. Pixels above this threshold were marked positive for APH-1α immunoreactivity and these data were divided by total number of pixels imaged to obtain relative pixels values across the different genotypes. Data were analyzed using one way ANOVA. A p value less than 0.05 was considered significant. All values are represented as SEM.

## Author Contributions and Notes

T.R.W., and J.L.E conceived the study, and developed the approach, T.R.W., P.A.G, and J.L. performed research, P.A.G and J.L. wrote software, T.R.W, P.A.G, D.M., and J.L.E. analyzed data; and T.R.W and J.L.E wrote the paper.

## Acknowledgments

P.A.G. supported by a training fellowship from the Gulf Coast Consortia, on the NLM Training Program in Biomedical Informatics & Data Science (T15LM007093).

## Notes

### Competing Interest Statement

The authors have declared no competing interest.

## References

1. Jellinger KA (2013) Pathology and pathogenesis of vascular cognitive impairment—a critical update. Front Aging Neurosci 5:. https://doi.org/10.3389/fnagi.2013.00017

2. Toledo JB, Arnold SE, Raible K, et al (2013) Contribution of cerebrovascular disease in autopsy confirmed neurodegenerative disease cases in the National Alzheimer’s Coordinating Centre. Brain J Neurol 136:2697–2706. https://doi.org/10.1093/brain/awt188

3. Attems J, Jellinger KA (2014) The overlap between vascular disease and Alzheimer’s disease--lessons from pathology. BMC Med 12:206. https://doi.org/10.1186/s12916-014-0206-2

4. Farkas E, Luiten PG (2001) Cerebral microvascular pathology in aging and Alzheimer’s disease. Prog Neurobiol 64:575–611. https://doi.org/10.1016/s0301-0082(00)00068-x

5. Savva GM, Wharton SB, Ince PG, et al (2009) Age, Neuropathology, and Dementia. N Engl J Med 360:2302–2309. https://doi.org/10.1056/NEJMoa0806142

6. Ribatti D (2014) History of research on angiogenesis. Chem Immunol Allergy 99:1–14. https://doi.org/10.1159/000353311

7. Imbimbo BP, Solfrizzi V, Panza F (2010) Are NSAIDs useful to treat Alzheimer’s disease or mild cognitive impairment? Front Aging Neurosci 2:. https://doi.org/10.3389/fnagi.2010.00019

8. McGeer PL (2000) Cyclo-oxygenase-2 inhibitors: rationale and therapeutic potential for Alzheimer’s disease. Drugs Aging 17:1–11. https://doi.org/10.2165/00002512-200017010-00001

9. Womack TR, Vollert C, Nwoko O, et al (2020) Prostacyclin Promotes Degenerative Pathology in a Model of Alzheimer’s disease. bioRxiv 2020.04.15.039842. https://doi.org/10.1101/2020.04.15.039842

10. Wang P, Guan P-P, Guo J-W, et al (2016) Prostaglandin I2 upregulates the expression of anterior pharynx-defective 1α and anterior pharynx-defective-1β in amyloid precursor protein/presenilin 1 transgenic mice. Aging Cell 15:861– 871. https://doi.org/10.1111/acel.12495

11. Arnaoutova I, George J, Kleinman HK, Benton G (2009) The endothelial cell tube formation assay on basement membrane turns 20: state of the science and the art. Angiogenesis 12:267–274. https://doi.org/10.1007/s10456-009-9146-4

12. De Strooper B (2003) Aph-1, Pen-2, and Nicastrin with Presenilin generate an active gamma-Secretase complex. Neuron 38:9–12. https://doi.org/10.1016/s0896-6273(03)00205-8

13. Stitham J, Midgett C, Martin K, Hwa J (2011) Prostacyclin: An Inflammatory Paradox. Front Pharmacol 2:. https://doi.org/10.3389/fphar.2011.00024

14. Freitas-Andrade M, Raman-Nair J, Lacoste B (2020) Structural and Functional Remodeling of the Brain Vasculature Following Stroke. Front Physiol 11:. https://doi.org/10.3389/fphys.2020.00948

15. Kawabe Jun-ichi, Yuhki Koh-ichi, Okada Motoi, et al (2010) Prostaglandin I2 Promotes Recruitment of Endothelial Progenitor Cells and Limits Vascular Remodeling. Arterioscler Thromb Vasc Biol 30:464–470. https://doi.org/10.1161/ATVBAHA.109.193730

16. Lambers C, Roth M, Jaksch P, et al (2018) Treprostinil inhibits proliferation and extracellular matrix deposition by fibroblasts through cAMP activation. Sci Rep 8:. https://doi.org/10.1038/s41598-018-19294-1

17. Schermuly RT, Yilmaz H, Ghofrani HA, et al (2005) Inhaled iloprost reverses vascular remodeling in chronic experimental pulmonary hypertension. Am J Respir Crit Care Med 172:358–363. https://doi.org/10.1164/rccm.200502-296OC

18. Pola R, Gaetani E, Flex A, et al (2004) Comparative analysis of the in vivo angiogenic properties of stable prostacyclin analogs: a possible role for peroxisome proliferator-activated receptors. J Mol Cell Cardiol 36:363– 370. https://doi.org/10.1016/j.yjmcc.2003.10.016

19. Wharton J, Davie N, Upton PD, et al (2000) Prostacyclin Analogues Differentially Inhibit Growth of Distal and Proximal Human Pulmonary Artery Smooth Muscle Cells. 7

20. Doganci S, Yildirim V, Yesildal F, et al (2015) Comparison of angiogenic and proliferative effects of three commonly used agents for pulmonary artery hypertension (sildenafil, iloprost, bosentan): is angiogenesis always beneficial? Eur Rev Med Pharmacol Sci 19:1900–1906

21. Hoang KG, Allison S, Murray M, Petrovic N (2015) Prostanoids regulate angiogenesis acting primarily on IP and EP4 receptors. Microvasc Res 101:127–134. https://doi.org/10.1016/j.mvr.2015.07.004

22. Farber HW (1991) Differences in Pulmonary and Systemic Arterial Endothelial Cell Adaptation to Chronic Hypoxia. In: Lahiri S, Cherniack NS, Fitzgerald RS (eds) Response and Adaptation to Hypoxia: Organ to Organelle. Springer, New York, NY, pp 195–201

23. Murata T, Ushikubi F, Matsuoka T, et al (1997) Altered pain perception and inflammatory response in mice lacking prostacyclin receptor. Nature 388:678–682. https://doi.org/10.1038/41780

24. Murohara Toyoaki, Horowitz Jeffrey R., Silver Marcy, et al (1998) Vascular Endothelial Growth Factor/Vascular Permeability Factor Enhances Vascular Permeability Via Nitric Oxide and Prostacyclin. Circulation 97:99–107. https://doi.org/10.1161/01.CIR.97.1.99

25. Muramatsu R, Kuroda M, Matoba K, et al (2015) Prostacyclin prevents pericyte loss and demyelination induced by lysophosphatidylcholine in the central nervous system. J Biol Chem 290:11515–11525. https://doi.org/10.1074/jbc.M114.587253

26. Abramovitz M, Adam M, Boie Y, et al (2000) The utilization of recombinant prostanoid receptors to determine the affinities and selectivities of prostaglandins and related analogs. Biochim Biophys Acta 1483:285–293. https://doi.org/10.1016/s1388-1981(99)00164-x

27. Whittle BJ, Silverstein AM, Mottola DM, Clapp LH (2012) Binding and activity of the prostacyclin receptor (IP) agonists, treprostinil and iloprost, at human prostanoid receptors: treprostinil is a potent DP1 and EP2 agonist. Biochem Pharmacol 84:68–75. https://doi.org/10.1016/j.bcp.2012.03.012

28. He T, Santhanam AVR, Lu T, et al (2017) Role of prostacyclin signaling in endothelial production of soluble amyloid precursor protein-α in cerebral microvessels. J Cereb Blood Flow Metab 37:106–122. https://doi.org/10.1177/0271678X15618977

29. Dries DR, Yu G (2008) ASSEMBLY, MATURATION, AND TRAFFICKING OF THE γ-SECRETASE COMPLEX IN ALZHEIMER’S DISEASE. Curr Alzheimer Res 5:132–146

30. Francis R, McGrath G, Zhang J, et al (2002) aph-1 and pen-2 are required for Notch pathway signaling, gamma-secretase cleavage of betaAPP, and presenilin protein accumulation. Dev Cell 3:85–97. https://doi.org/10.1016/s1534-5807(02)00189-2

31. Carroll CM, Li Y-M (2016) Physiological and pathological roles of the γ-secretase complex. Brain Res Bull 126:199–206. https://doi.org/10.1016/j.brainresbull.2016.04.019

32. Rius-Pérez S, Tormos AM, Pérez S, Taléns-Visconti R (2018) Vascular pathology: Cause or effect in Alzheimer disease? Neurol Engl Ed 33:112–120. https://doi.org/10.1016/j.nrleng.2015.07.008

33. Dunn KW, Kamocka MM, McDonald JH (2011) A practical guide to evaluating colocalization in biological microscopy. Am J Physiol-Cell Physiol 300:C723–C742. https://doi.org/10.1152/ajpcell.00462.2010

34. Li J, Artur C, Eriksen J, et al (2020) Segmenting Continuous but Sparsely-Labeled Structures in Super-Resolution Microscopy Using Perceptual Grouping. In: Medical Image Computing and Computer Assisted Intervention – MICCAI 2020. Springer, Cham, pp 141–150

